## Supplemental Files for "Conformational trajectory of the HIV-1 fusion peptide during CD4-induced envelope opening"

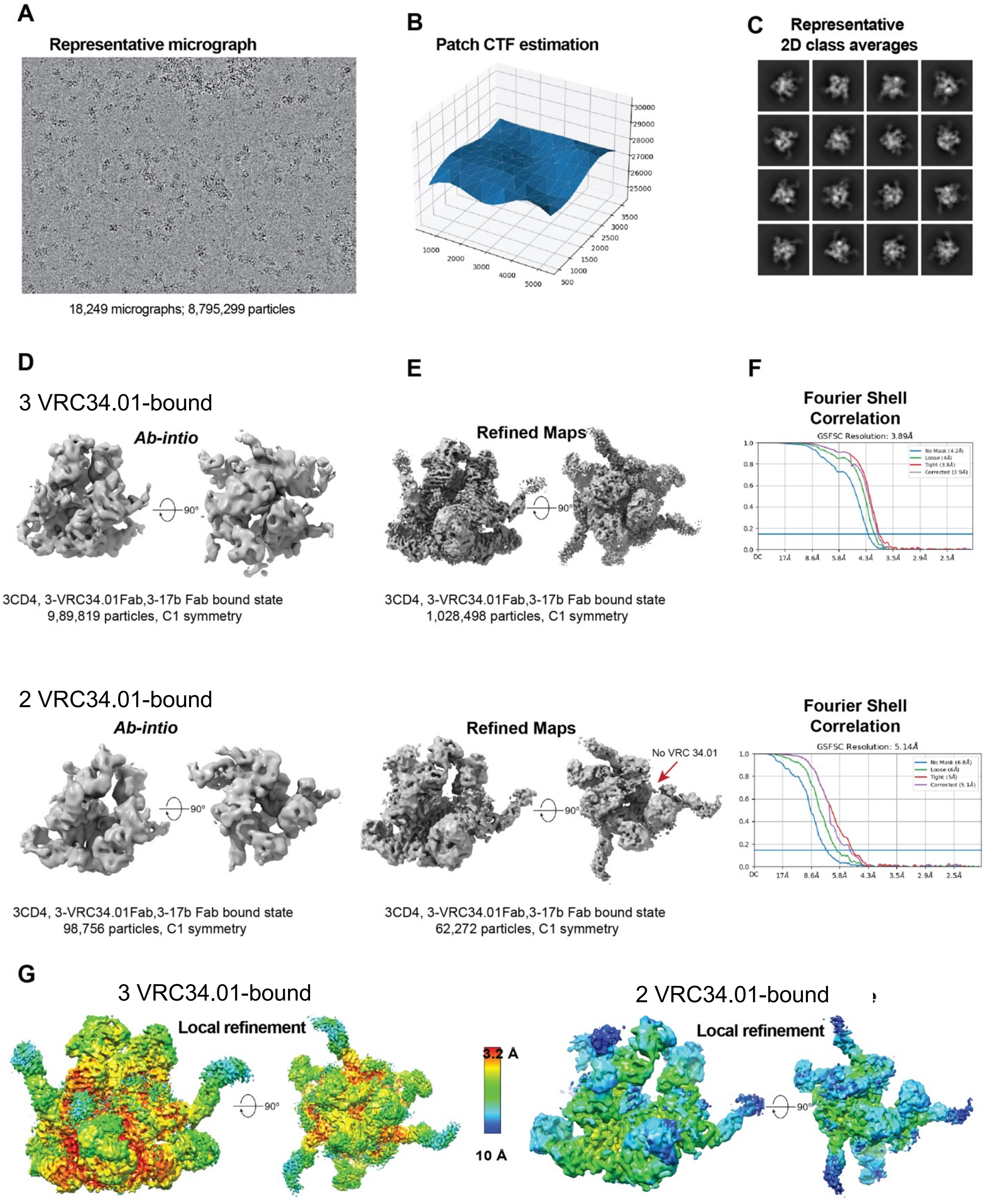
A
Representative micrograph
18,249 micrographs; 8,795,299 particles
B
Patch CTF estimation
C
Representative
2D class averages
D
3 VRC34.01-bound
Ab-intio
90°
3CD4, 3-VRC34.01Fab,3-17b Fab bound state
9,89,819 particles, C1 symmetry
E
Refined Maps
3CD4, 3-VRC34.01Fab,3-17b Fab bound state
1,028,498 particles, C1 symmetry
F
Fourier Shell Correlation
GSFSC Resolution: 3.89Å
2 VRC34.01-bound
Ab-intio
3CD4, 3-VRC34.01Fab,3-17b Fab bound state
98,756 particles, C1 symmetry
Refined Maps
No VRC 34.01
3CD4, 3-VRC34.01Fab,3-17b Fab bound state
62,272 particles, C1 symmetry
Fourier Shell Correlation
GSFSC Resolution: 5.14Å
G
3 VRC34.01-bound
Local refinement
3.2 Å
10 Å
2 VRC34.01-bound
Local refinement

**Figure S1. Cryo-EM data processing for BG505 SOSIP.664 incubated with CD4 and 17b for 1.3 hours followed by incubation with VRC34.01 Fab for 30 minutes prior to vitrification**. **(A)** Representative frame aligned micrograph. (B) Defocus value estimation across micrograph during Patch CTF estimation. **(C)** Representative 2D class averages. Particle extraction box size = 345.6 Å. **(D)** *Ab intio* 3D reconstructions of CD4,17b-bound Env complexes with either three or two VRC34.01 Fabs bound per Env trimer. **(E)** Refined maps for corresponding states shown in panel **D**. **(F)** Fourier shell correlation (FSC) curves for refined 3D maps shown in panel **E** with the horizontal blue line showing FSC threshold value of 0.143. **(G)** Refined maps colored by local resolution.

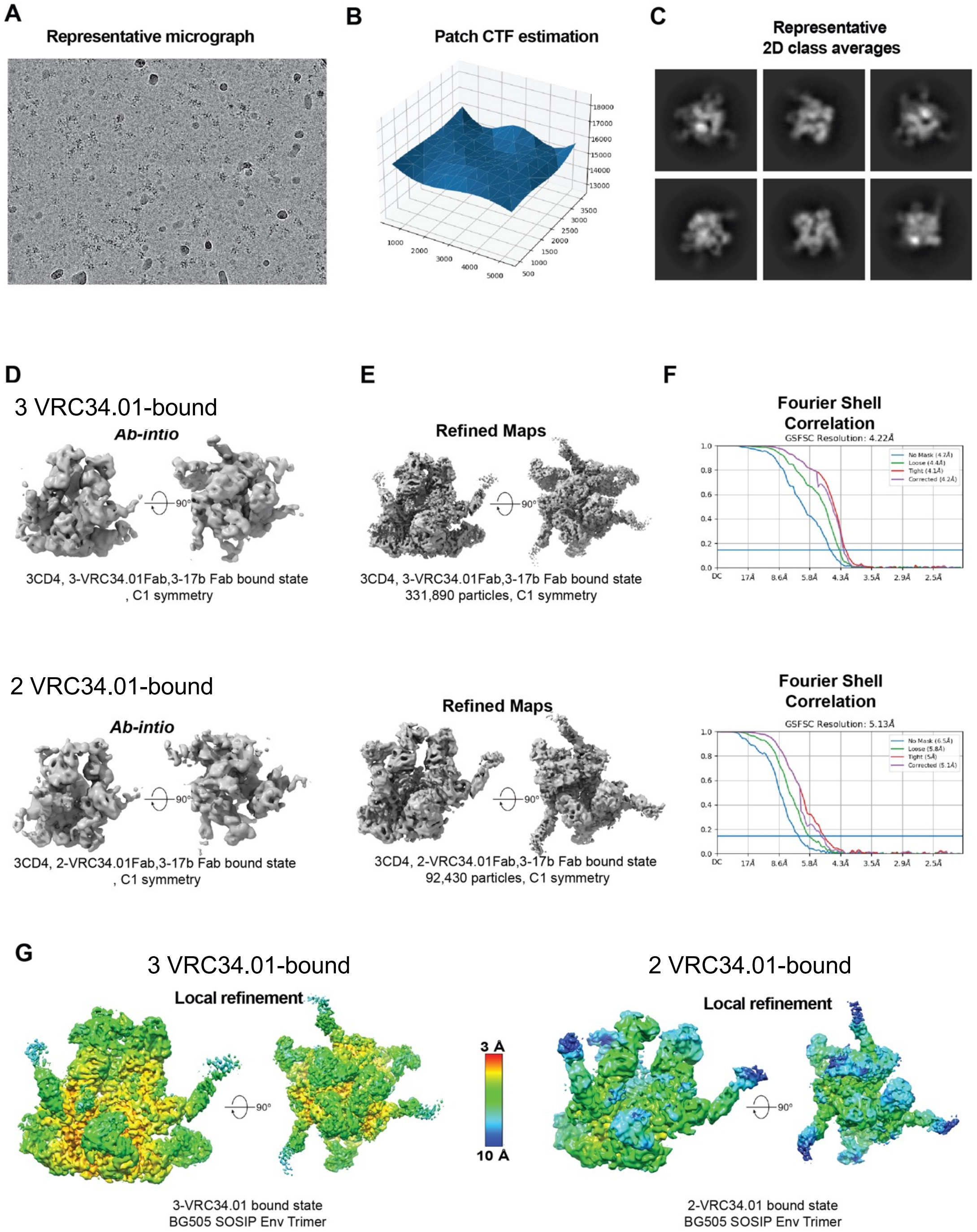

A
Representative micrograph
B
Patch CTF estimation
C
Representative
2D class averages
D
3 VRC34.01-bound
Ab-intio
90°
3CD4, 3-VRC34.01Fab,3-17b Fab bound state
, C1 symmetry
E
Refined Maps
90°
3CD4, 3-VRC34.01Fab,3-17b Fab bound state
331,890 particles, C1 symmetry
F
Fourier Shell
Correlation
GSFSC Resolution: 4.22Å
No Mask (4.7Å)
Loose (4.4Å)
Tight (4.1Å)
Corrected (4.2Å)
2 VRC34.01-bound
Ab-intio
90°
3CD4, 2-VRC34.01Fab,3-17b Fab bound state
, C1 symmetry
Refined Maps
90°
3CD4, 2-VRC34.01Fab,3-17b Fab bound state
92,430 particles, C1 symmetry
Fourier Shell
Correlation
GSFSC Resolution: 5.13Å
No Mask (6.5Å)
Loose (5.8Å)
Tight (5Å)
Corrected (5.1Å)
G
3 VRC34.01-bound
Local refinement
90°
3-VRC34.01 bound state
BG505 SOSIP Env Trimer
3 Å
10 Å
2 VRC34.01-bound
Local refinement
90°
2-VRC34.01 bound state
BG505 SOSIP Env Trimer

**Figure S2. Cryo-EM data processing for BG505 SOSIP.664 incubated with CD4 and 17b for 20 hours followed by incubation with VRC34.01 Fab for 30 minutes prior to vitrification**. **(A)** Representative frame aligned micrograph. (B) Defocus value estimation across micrograph during Patch CTF estimation. **(C)** Representative 2D class averages. Particle extraction box size = 345.6 Å. **(D)** *Ab intio* 3D reconstructions of CD4,17b-bound Env complexes with either three or two VRC34.01 Fabs bound per Env trimer. **(E)** Refined maps for corresponding states shown in panel **D**. **(F)** Fourier shell correlation (FSC) curves for refined 3D maps shown in panel **E** with the horizontal blue line showing FSC threshold value of 0.143. **(G)** Refined maps colored by local resolution.

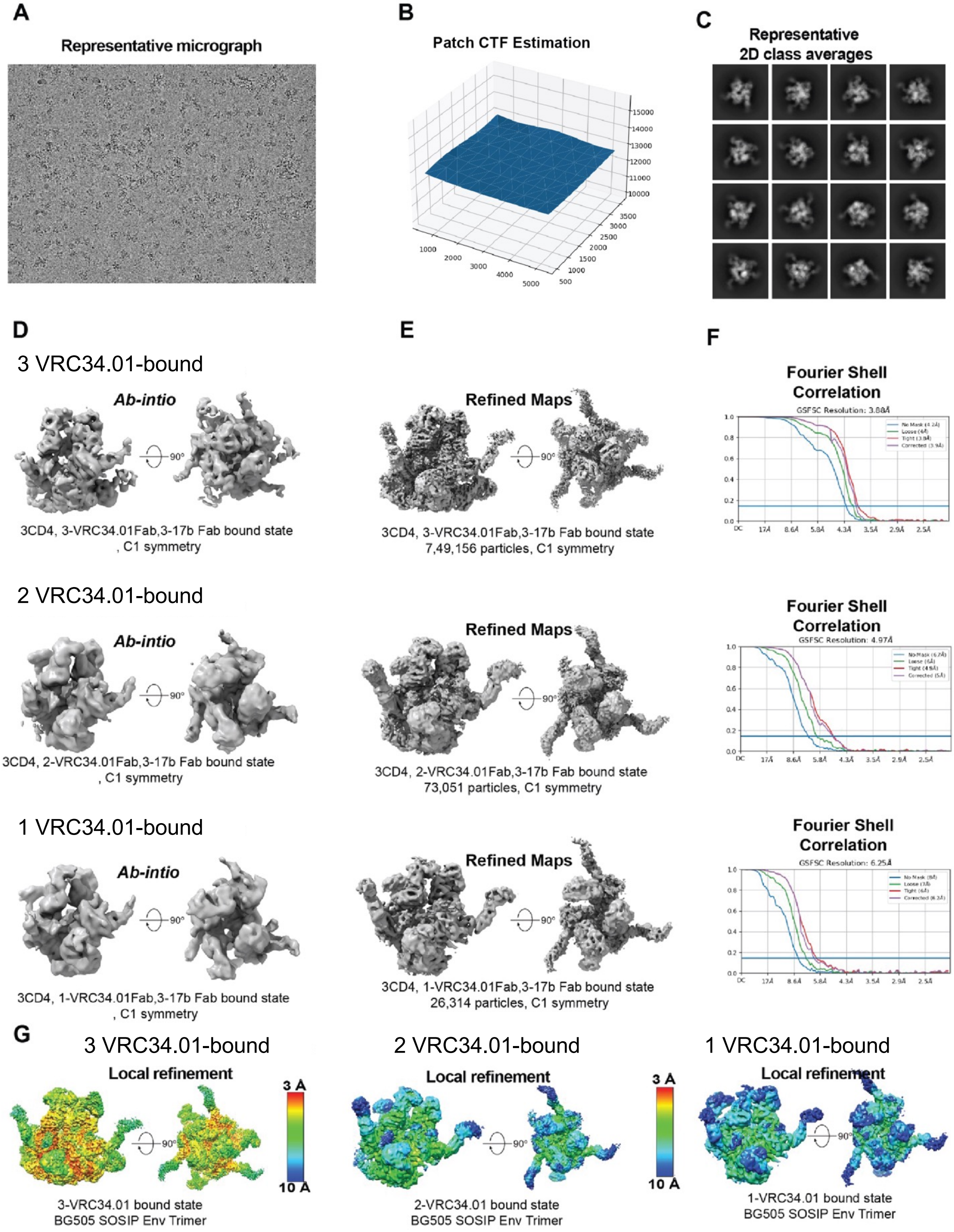
A
Representative micrograph
B
Patch CTF Estimation
15000
14000
13000
12000
11000
10000
1000
2000
3000
4000
5000
3500
3000
2500
2000
1500
1000
500
C
Representative
2D class averages
D
3 VRC34.01-bound
Ab-intio
90°
3CD4, 3-VRC34.01Fab,3-17b Fab bound state
, C1 symmetry
E
Refined Maps
90°
3CD4, 3-VRC34.01Fab,3-17b Fab bound state
7,49,156 particles, C1 symmetry
F
Fourier Shell
Correlation
GSFSC Resolution: 3.88Å
2 VRC34.01-bound
Ab-intio
90°
3CD4, 2-VRC34.01Fab,3-17b Fab bound state
, C1 symmetry
Refined Maps
90°
3CD4, 2-VRC34.01Fab,3-17b Fab bound state
73,051 particles, C1 symmetry
Fourier Shell
Correlation
GSFSC Resolution: 4.97Å
1 VRC34.01-bound
Ab-intio
90°
3CD4, 1-VRC34.01Fab,3-17b Fab bound state
, C1 symmetry
Refined Maps
90°
3CD4, 1-VRC34.01Fab,3-17b Fab bound state
26,314 particles, C1 symmetry
Fourier Shell
Correlation
GSFSC Resolution: 6.25Å
G
3 VRC34.01-bound
Local refinement
90°
3 Å
10 Å
3-VRC34.01 bound state
BG505 SOSIP Env Trimer
2 VRC34.01-bound
Local refinement
90°
3 Å
10 Å
2-VRC34.01 bound state
BG505 SOSIP Env Trimer
1 VRC34.01-bound
Local refinement
90°
1-VRC34.01 bound state
BG505 SOSIP Env Trimer

**Figure S3. Cryo-EM data processing for BG505 SOSIP.664 incubated with CD4 and 17b for 3 days followed by incubation with VRC34.01 Fab for 30 minutes prior to vitrification**. **(A)** Representative frame aligned micrograph. (B) Defocus value estimation across micrograph during Patch CTF estimation. **(C)** Representative 2D class averages. Particle extraction box size = 345.6 Å. **(D)** *Ab intio* 3D reconstructions of CD4,17b-bound Env complexes with either three, two or one VRC34.01 Fabs bound per Env trimer. **(E)** Refined maps for corresponding states shown in panel **D**. **(F)** Fourier shell correlation (FSC) curves for refined 3D maps shown in panel **E** with the horizontal blue line showing FSC threshold value of 0.143. **(G)** Refined maps colored by local resolution.

**A**

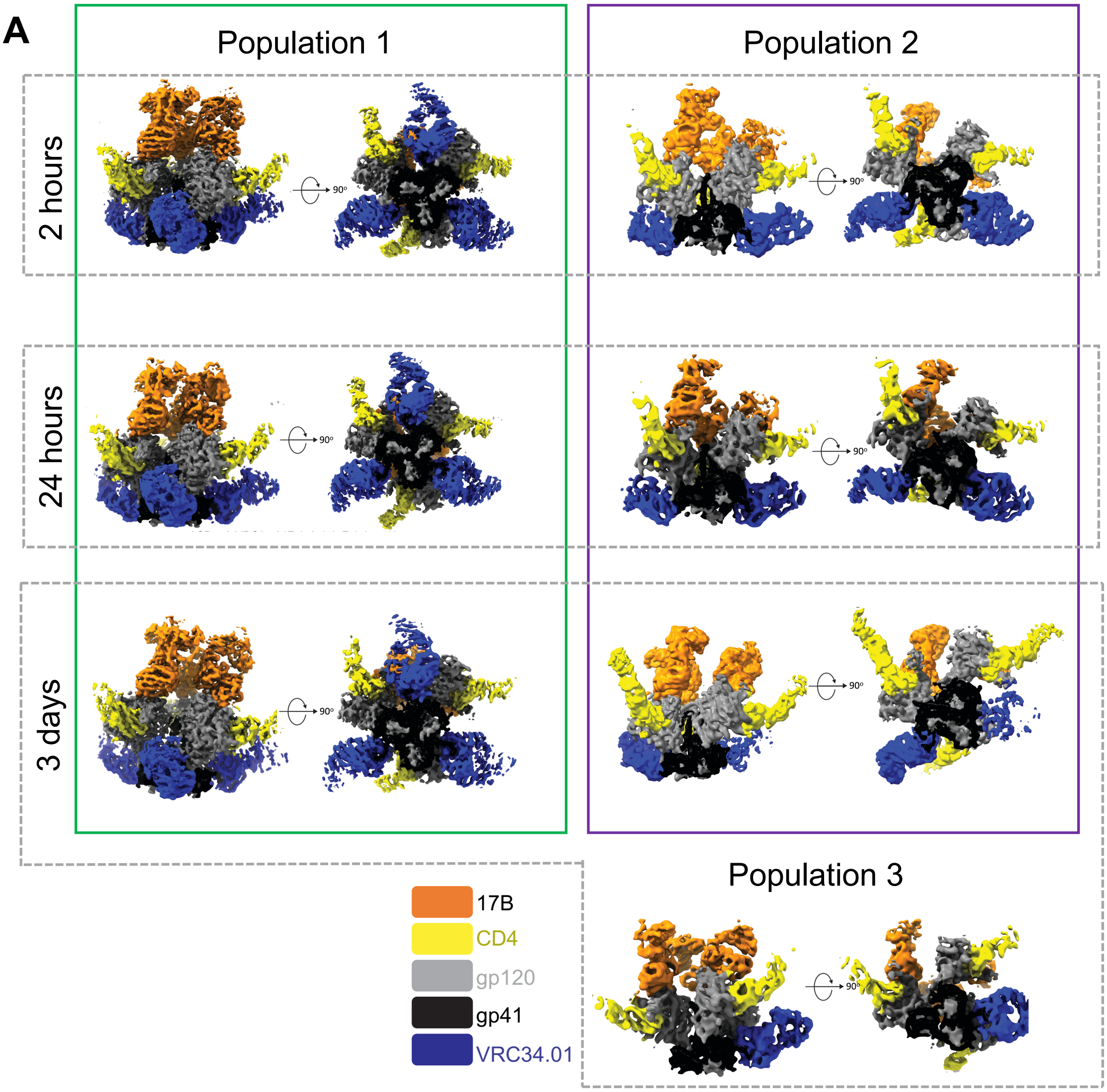


**B**

| | 3-CD4, 3-17b Fab, 3-VRC34.01 Fab bound **Population 1** | 3-CD4, 3-17b Fab, 2-VRC34.01 Fab bound **Population 2** | 3-CD4, 3-17b Fab, 2-VRC34.01 Fab bound **Population 3** |
|---|---|---|---|
| 13 hours | 1,028,496 (94.3 %) | 62,272 (5.7 %) | ND |
| 20 hours | 331,890 (78.2 %) | 92,430 (21.8 %) | ND |
| 3 days | 749,156 (88.3 %) | 73,051 (8.6 %) | 26,314 (3.1 %) |

**Figure S4. Diversity of HIV-1 Env intermediate states visualized by cryo-EM. (A)** 3D reconstructions of particle populations identified in cryo-EM datasets at different time-points post CD4, 17b incubation of HIV-1 Env followed by addition of VRC34.01 Fab 30 minutes prior to sample vitrification. **(B)** Summary of approximate particle numbers and relative percentages of each population identified at each timepoint.

**A** BG505 SOSIP-PGT122-VRC34.01 (Closed conformation)
PDB: 5I8H

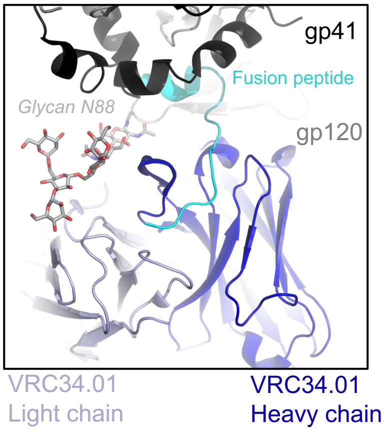


**B** BG505 SOSIP-CD4-17b-VRC34.01 (Partially open conformation)
PDB: 9D90 (this study)

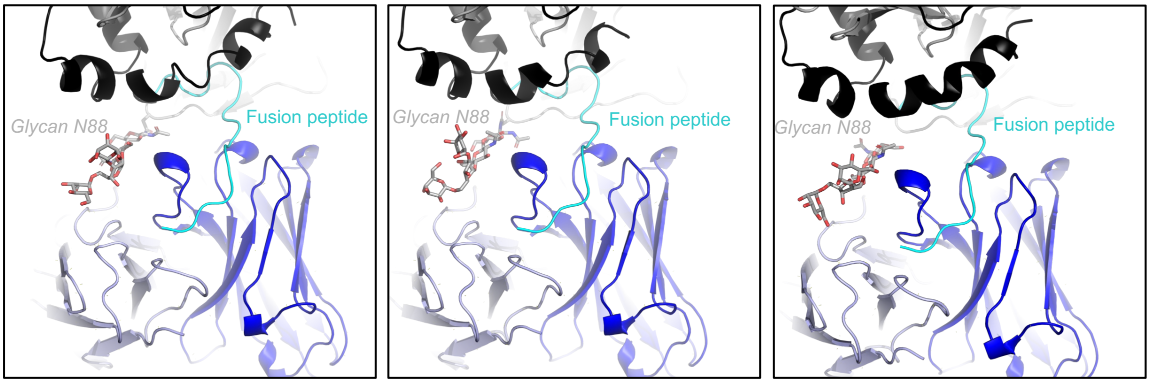


**C** BG505 SOSIP-CD4-17b-VRC34.01 (Partially open conformation)
PDB: 9D90 (this study)

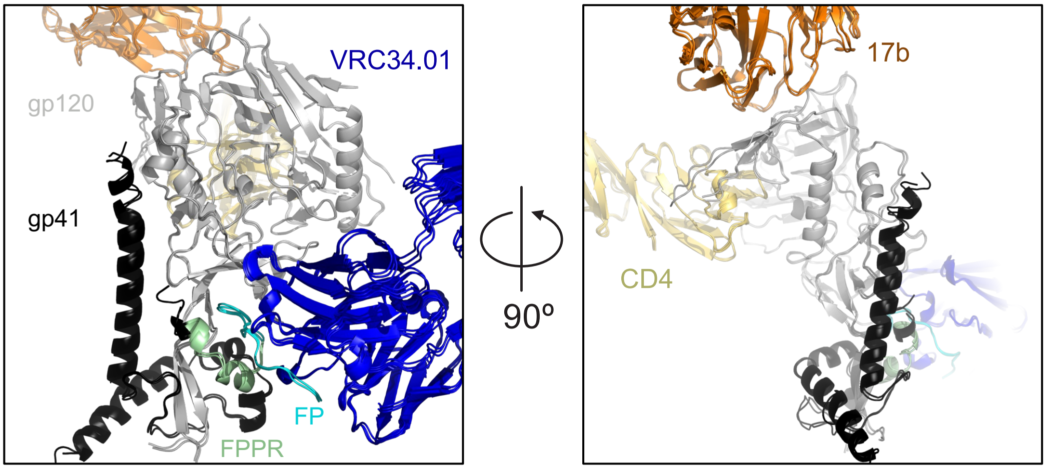

**Figure S5. Antibody recognition of FP in the context of diverse Env conformations. (A)** VRC34.01 bound to HIV-1 Env in closed conformation. VRC34.01 heavy and light chains are shown as dark blue and light blue cartoon, respectively; gp120 is colored grey with the glycan at position N88 shown in stock representation; gp41 is colored black; FP s colored cyan. **(B)** VRC34.01 bound to HIV-1 Env in partially open conformation. Color scheme is same as in panel **A**. **(C)** The three protomers of the Population 1 structure (PDB: 9D90; this study) overlayed by their gp120 subunits. VRC34.01 bound to HIV-1 Env in partially open conformation. Color scheme is same as in panel **A,** except both VRC34.01 heavy and light chains are colored dark blue. FPPR is colored pale green, CD4 yellow and 17b orange.

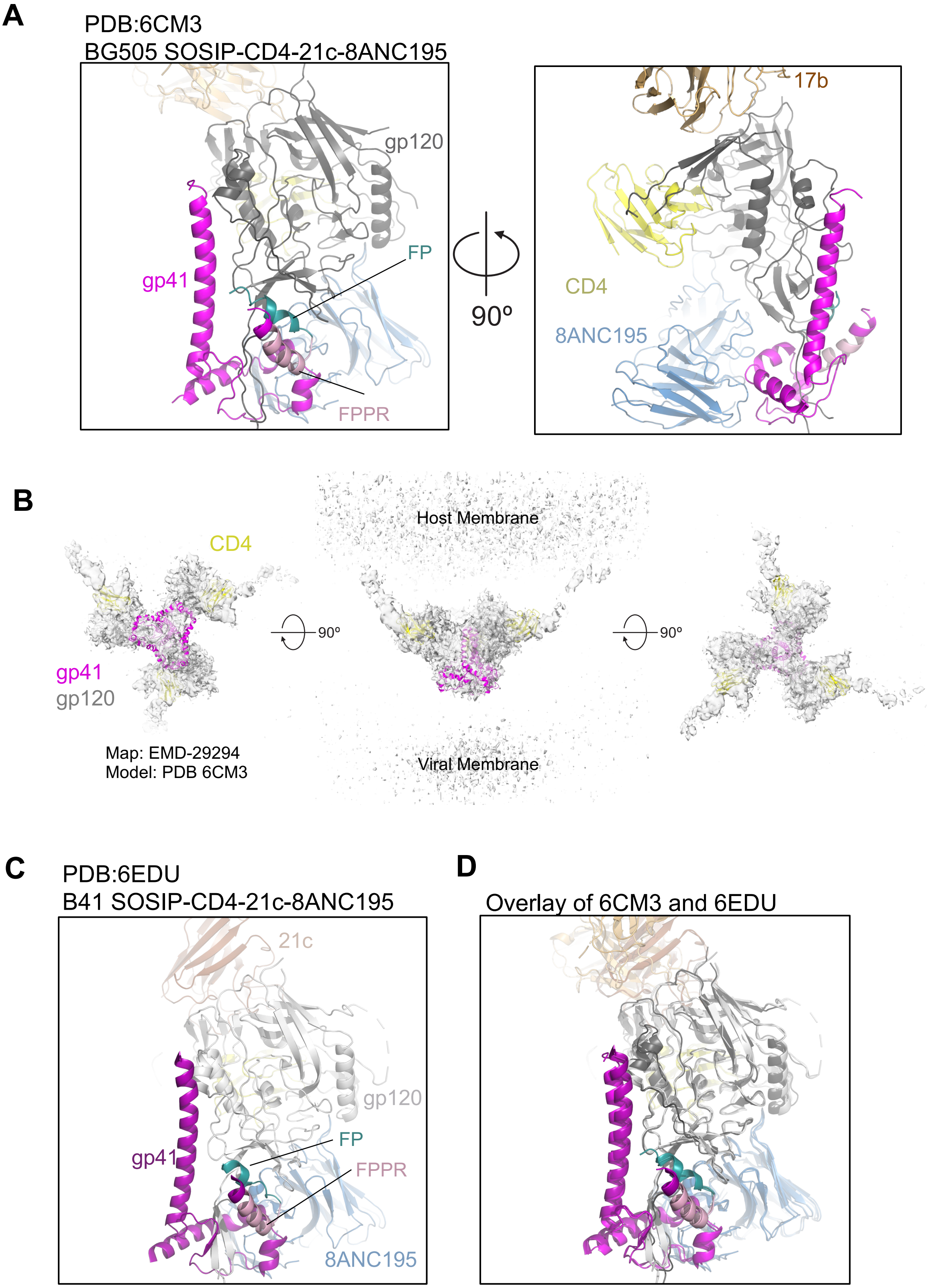

A
PDB:6CM3
BG505 SOSIP-CD4-21c-8ANC195
gp120
FP
gp41
FPPR
90º
17b
CD4
8ANC195
B
Host Membrane
CD4
90º
90º
gp41
gp120
Map: EMD-29294
Model: PDB 6CM3
Viral Membrane
C
PDB:6EDU
B41 SOSIP-CD4-21c-8ANC195
21c
gp120
FP
gp41
FPPR
8ANC195
D
Overlay of 6CM3 and 6EDU

**Figure S6. Cryo-EM structure of BG505 SOSIP Env bound to CD4, 17b and 8ANC195 (PDB: 6CM3)**. **(A)** One protomer of the partially open Env bound to CD4, 17b Fab and 8ANC195 Fab (PDB ID: 6CM3) shown in cartoon representation zoomed-in at the gp120/gp41 interface. The gp120 subunit is colored dark grey, gp41 magenta, FP dark teal and FPPR light pink. The bound CD4 is colored yellow, 8ANC195 Fab marine blue, 17b Fab brown. **(B)** Coordinates of PDB: 6CM3 including Env (gp120 in light gray, gp41 in magenta) and CD4 (yellow) fitted into the *in situ* cryo-ET reconstruction of a partially open CD4-bound Env (EMD-29294). **(C)** One protomer of the partially open Env bound to CD4, 21c Fab and 8ANC195 Fab (PDB ID: 6EDU) shown in cartoon representation zoomed-in at the gp120/gp41 interface. The gp120 subunit is colored light grey, gp41 dark purple, FP dark teal and FPPR light pink. The bound CD4 is colored yellow, 8ANC195 Fab marine blue, 21c Fab chocolate. **(D)** Structures of BG505 SOSIP bound to CD4, 17b Fab and 8ANC195 Fab, and of B41 SOSIP bound to CD4, 21c Fab and 8ANC195 Fab, overlaid by their gp120 subunits.

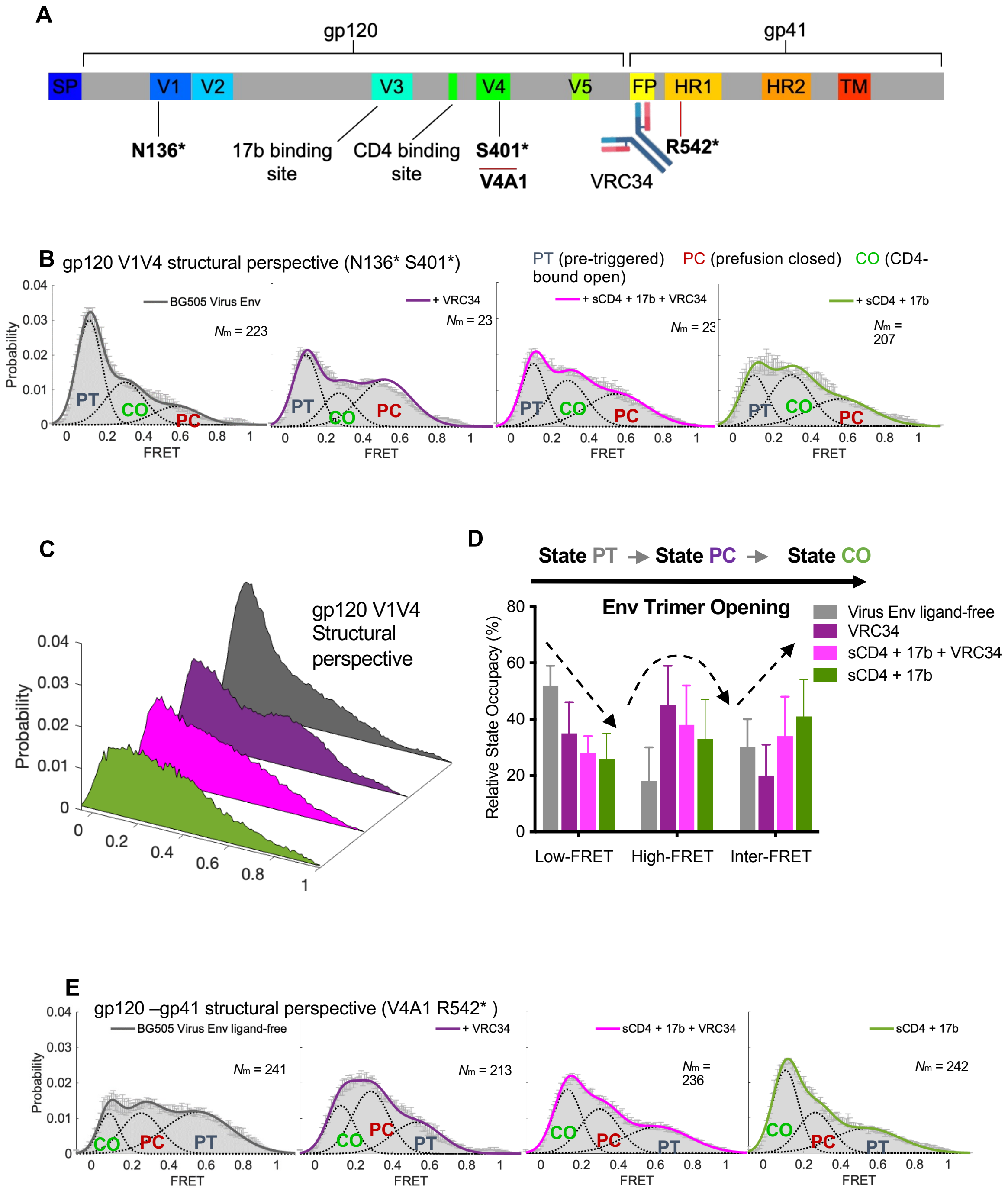
A
gp120
gp41
SP
V1
V2
V3
V4
V5
FP
HR1
HR2
TM
N136*
17b binding site
CD4 binding site
S401*
V4A1
VRC34
R542*
B
gp120 V1V4 structural perspective (N136* S401*)
PT (pre-triggered) PC (prefusion closed) CO (CD4-bound open)
BG505 Virus Env
+ VRC34
+ sCD4 + 17b + VRC34
+ sCD4 + 17b
Probability
FRET
PT
CO
PC
C
gp120 V1V4 Structural perspective
Probability
D
State PT → State PC → State CO
Env Trimer Opening
Relative State Occupacy (%)
Virus Env ligand-free
VRC34
sCD4 + 17b + VRC34
sCD4 + 17b
Low-FRET
High-FRET
Inter-FRET
E
gp120 –gp41 structural perspective (V4A1 R542* )
BG505 Virus Env ligand-free
+ VRC34
sCD4 + 17b + VRC34
sCD4 + 17b
Probability
FRET
CO
PC
PT

**Fig S7. Visualizing VRC34-related conformational stabilization and activation of virus Env by smFRET of full-length Env on native virions from two structural perspectives.**

(**A**) The domain organization of $Env_{BG505}$ sequence is depicted, with the fluorescent labeling sites used for smFRET and antibody recognition clearly indicated. FRET-paired Cy3/Cy5 fluorophores were site-specifically labeled on V1V4 (N136* S401*) or gp120-gp41 (V4A1 R542*). A1 is a peptide tag, and *indicates an unnatural amino acid incorporation site for click labeling.

(**B**) The conformational distribution-indicated FRET histograms of virus $Env_{BG505}$ probed from gp120 V1V4 (N136* S401*) structural perspectives in the absence or presence of different ligands. Virus $Env_{BG505}$ samples three primary conformational states (PT: Pre-triggered, PC: Pre-receptor Closed, and CO: CD4-bound open). PT predominates in the ligand-free condition (left panel), while VRC34 (two middle panels) shifts the conformational landscape differently from that of the CD4-bound opening (right panel). FRET histogram represents the mean + s.e.m. determined from three randomly assigned populations of FRET traces ($N_m$, number of traces).

(**C**) Three-dimensional presentation and **(D**) state occupancy quantifications of FRET histograms in B of virus $Env_{BG505}$ from the gp120 V1V4 perspective. Relative state occupancies are presented as mean ± s.e.m. determined by estimating the area under each Gaussian curve of histograms in B. For state occupancies and parameters, see Table S2 and Methods.

**(E**) The conformational distribution-indicated FRET histograms of virus $Env_{BG505}$ probed from the gp120 –gp41 (V4A1 R542*) structural perspectives in the absence or presence of different ligands. Virus $Env_{BG505}$ samples three primary conformational states (PT: Pre-triggered, PC: Pre-receptor Closed, and CO: CD4-bound open). PT predominates in the ligand-free condition (left panel), while VRC34 (two middle panels) shifts the conformational landscape differently from that of the CD4-bound opening (right panel). FRET histogram represents the mean + s.e.m. determined from three randomly assigned populations of FRET traces ($N_m$, number of traces). Three-dimensional presentation and quantification of FRET histograms in from the gp120-gp41 perspective and shown in **Figure 4E.**

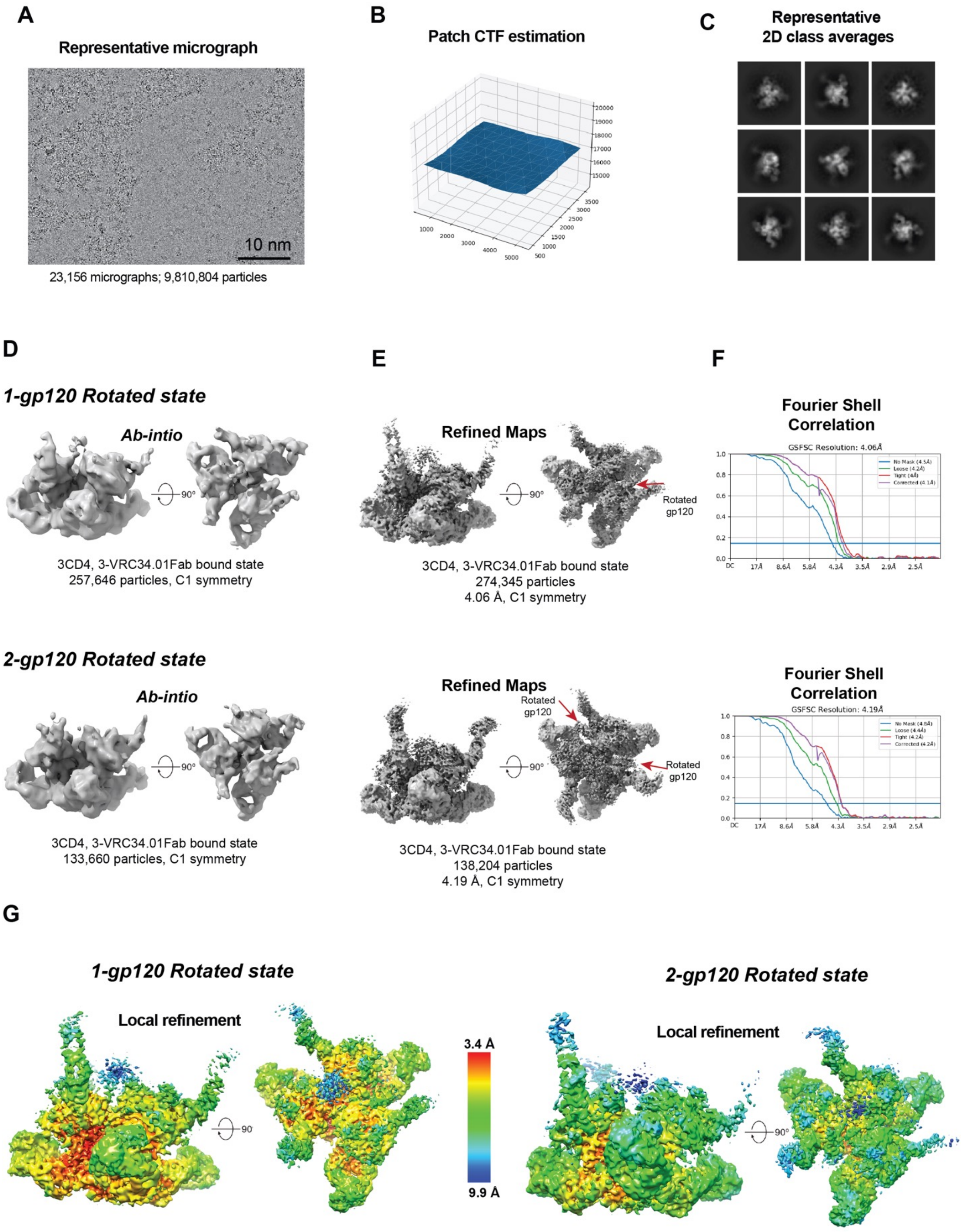
A
Representative micrograph
10 nm
23,156 micrographs; 9,810,804 particles
B
Patch CTF estimation
C
Representative
2D class averages
D
1-gp120 Rotated state
Ab-intio
90°
3CD4, 3-VRC34.01Fab bound state
257,646 particles, C1 symmetry
E
Refined Maps
90°
Rotated
gp120
3CD4, 3-VRC34.01Fab bound state
274,345 particles
4.06 Å, C1 symmetry
F
Fourier Shell
Correlation
GSFSC Resolution: 4.06Å
No Mask (4.5Å)
Loose (4.2Å)
Tight (4Å)
Corrected (4.1Å)
DC
17Å
8.6Å
5.8Å
4.3Å
3.5Å
2.9Å
2.5Å
2-gp120 Rotated state
Ab-intio
90°
3CD4, 3-VRC34.01Fab bound state
133,660 particles, C1 symmetry
Refined Maps
Rotated
gp120
90°
Rotated
gp120
3CD4, 3-VRC34.01Fab bound state
138,204 particles
4.19 Å, C1 symmetry
Fourier Shell
Correlation
GSFSC Resolution: 4.19Å
No Mask (4.8Å)
Loose (4.4Å)
Tight (4.2Å)
Corrected (4.2Å)
G
1-gp120 Rotated state
Local refinement
3.4 Å
9.9 Å
2-gp120 Rotated state
Local refinement
90°

**Figure S8. Cryo-EM data processing for BG 505 SOSIP.664 gp120 incubated with CD4 for 1.3 HR followed by incubation for VRC34.01 for 30 minutes**. **(A)** Representative frame aligned micrograph. (B) Defocus value estimation across micrograph during PATCH CTF estimation. **(C)** Representative 2D class average of extracted particles showing particles in different orientations. Particle extraction box size 345.6 Å **(D)** Ab intio 3D reconstruction dividing particles in one and two gp120 rotated states. **(E)** Refined maps for corresponding states shown in D. **(F)** Fourier shell correlation (FCS) curves for refined 3D maps shown in figure E with the horizontal blue line showing FCS threshold value of 0.143. **(G)** Refined maps color coded by local resolution.

| BG505 Virus Env | Curve fitting $R^2$ | RMSE | Pre-triggered | Pre-fusion closed | Three-CD4 open |
|---|---|---|---|---|---|
| | | | μ: 0.1 ± 0.02 | μ: 0.60 ± 0.03 | μ: 0.3 ± 0.03 |
| | | | σ: 0.07 ± 0.01 | σ: 0.15 ± 0.02 | σ: 0.1 ± 0.01 |
| **$N136_{TAG}$ $S401_{TAG}$** | | | | | |
| Ligand-free | 0.9891 | 9.4074e-04 | 52% ± 07% | 18% ± 12% | 30% ± 10% |
| + VRC34 | 0.9669 | 0.0012 | 35% ± 11% | 45% ± 14% | 20% ± 11% |
| + sCD4 + 17b + VRC34 | 0.9891 | 7.044e-04 | 28% ± 06% | 38% ± 14% | 34% ± 14% |
| + sCD4 + 17b | 0.9791 | 9.2681e-04 | 26% ± 09% | 33% ± 14% | 41% ± 13% |

| BG505 Virus Env | Curve fitting $R^2$ | RMSE | | | |
|---|---|---|---|---|---|
| | | | μ: 0.55 ± 0.05 | μ: 0.28 ± 0.03 | μ: 0.10 ± 0.02 |
| | | | σ: 0.15 ± 0.02 | σ: 0.1 ± 0.01 | σ: 0.07 ± 0.01 |
| **V4-A1 $R542_{TAG}$** | | | | | |
| Ligand-free | 0.9914 | 5.0798e-04 | 53% ± 07% | 30% ± 07% | 17% ± 03% |
| + VRC34 | 0.9903 | 7.6231e-04 | 36% ± 16% | 47% ± 12% | 17% ± 13% |
| + sCD4 + 17b + VRC34 | 0.9946 | 4.9572e-04 | 33% ± 06% | 33% ± 11% | 34% ± 07% |
| + sCD4 + 17b | 0.9914 | 7.3402e-04 | 32% ± 10% | 27% ± 14% | 41% ± 10% |

**Table S2. Relative state occupancy and fitting parameters of conformational distributions of the virus Env observed from two different structural perspectives, with reference to Figure 4E.**

A pair of fluorophores labeled on different Env sites result in different FRET signals. In each FRET structural perspective, FRET histograms were fitted into the sum of three distinct Gaussian distributions with defined means (μ) and deviations (σ) of the individual Gaussian/normal distributions. μ and σ were determined based on the observation of unbiased raw FRET signals and were further determined using hidden Markov modeling. As a measure of uncertainty, we presented the probability of each state Env occupies as mean + s.e.m. $R^2$ and RMSE (Root Mean Square Deviation) evaluate the goodness of fitting.
